# Assessment of the pollution incident performance of water and sewerage companies in England

**DOI:** 10.1101/2021.04.21.440747

**Authors:** Sarah Purnell, Nick Mills, Keith Davis, Christopher Joyce

**Author notes:** Corresponding author; address: University of Brighton, Cockcroft Building, Lewes Road, Brighton, BN2 4GJ, United Kingdom.

## Abstract

Comparison of water and sewerage company pollution performance in relation to severity, frequency and self-reporting of pollution incidents is made difficult by differences in environmental and operational conditions. In England, the deterioration in pollution incident performance across the water industry, makes it important to investigate common trends that could be addressed at national and regional levels. Yet, to date there has been no external peer-reviewed analysis of national pollution incident data in England. This project aimed to analyse available pollution incident data to assess and compare the performance of water and sewerage companies in England. Results indicated that there was significant variation in pollution incident numbers and the severity of the impact on the water environment for different asset types (operational property of the water and sewerage company such as a sewage treatment works). Increasing numbers of pollution incidents from pumping stations and sewage treatment works, were largely responsible for overall increases in pollution incidents. The highest increase in pollution incidents in 2019, was observed from pumping stations. Variation was evident in company self-reporting percentages across asset types. There were significant positive relationships between the self-reporting percentages of pollution incidents and total numbers of reported pollution incidents per 10,000 km sewer length for pumping stations and sewage treatment works. These results indicate that in at least these asset types, an estimated 5% of pollution incidents could go unreported, if not self-reported by the company. Pollution events not reported quickly by companies, can lead to more severe impacts to the water environment so rapid and consistent reporting of incidents is crucial for limiting damage. The results have significance for the water industry internationally, because the issues presented here are not restricted to England. Whilst this research highlighted a number of key areas for more detailed analysis, in the short-term, research should focus on investigating best practice for reporting pollution incidents. It is important to get an accurate baseline of the number of pollution incidents and whether a proportion are currently going unreported. This research should seek to aid the standardisation of reporting practice across the water industry.

## Introduction

In 2000, the EU Water Framework Directive (WFD) set out ambitious objectives to achieve good ecological and chemical status in all water bodies in the EU. In the UK, after leaving the EU, the WFD has been transposed into legislation (Water Environment Regulations, 2017). Achieving the objectives of the WFD has proved challenging and in 2015, only 47% of surface waters had good ecological status across the EU (EU Commission 2012; Voulvoulis et al., 2017). In the UK, WFD programmes of measures have not resulted in improved overall status of rivers. In the most recent WFD classification, only 16% of surface waters were classified as achieving good ecological status (EA, 2020a). Figures released in 2020, also showed that not a single river in England achieved a good chemical status (EA, 2020a). In 2019, the nine water and sewerage companies (WASC) operating in England reported 2192 pollution incidents that negatively impacted the water environment. From these incidents, 88% were listed as sewage materials. The introduction of contaminants found in sewage into aquatic environments leads to ecological deterioration of water bodies and presents a risk to human health (Kidd et al., 2007; Emmanuel et al., 2007; Cunningham et al., 2010; Rivera-Jaimes et al., 2018; Li et al., 2020; Purnell et al., 2020). Sewage contains high organic loads, nutrients and a large range of other contaminants including pharmaceuticals, personal care products, endocrine disrupting chemicals, metals, microplastics and pathogenic microorganisms (Köck-Schulmeyer et al., 2011; Al Aukidy and Verlicchi, 2012; Li et al., 2018; Purnell et al., 2016; Tran et al., 2019). Serious pollution incidents can result in large scale aquatic organism mortality events, but frequent low-level chronic exposure to contaminants can be equally damaging in the long term (Saaristo et al., 2018). Therefore, improving WASC pollution performance plays a key role in improving ecological and chemical status in water bodies.

The English Environment Agency (EEA), first introduced the Environmental Performance Assessment (EPA) in 2011, the aim of which is to assess and compare the environmental performance of the nine WASCs in England, annually (EA, 2019a). Within the EPA, environmental performance is quantitatively assessed by a series of six performance indicators: 1) total pollution incidents, 2) serious pollution incidents, 3) compliance with discharge permits, 4) self-reporting of pollution incidents, 5) delivery of environmental improvement schemes and 6) the provision of secure supplies of water (EA, 2019a). The EEA define an incident as “*a specific event or occurrence brought to the attention of the Environment Agency, within their areas of responsibility, which may have an environmental and/or operational impact*” (EA, 2016a). The EEA (2016a) assign pollution incidents to one of the following environmental impact categories:

- Category 1 – major, serious, persistent and/or extensive impact or effect on the environment, people and/or property.
- Category 2 – significant impact or effect on the environment, people and/or property.
- Category 3 – minor or minimal impact or effect on the environment, people and/or property.
- Category 4 – substantiated incident with no impact.

Since the introduction of the EPA in 2011, the EEA has used these numerical environmental performance indicators to compare the performance of the nine WASCs in England (EA, 2017a). The EPA is a non-statutory tool, that forms part of a wider assessment, facilitated through ongoing engagement and review meetings between water and sewerage companies (WASCs) and the EEA (EA, 2017a). Environmental performance comparisons aim to hold water and sewerage companies to account and drive improvement within the industry. Of the numerical environmental performance indicators reported in annual EPAs, three have relevance to this research; category 1-3 pollution incidents per 10,000 km, serious category 1-2 pollution incidents per 10,000km and the percentage of self-reported pollution incidents. To aid clarity and to keep text concise, reference to ‘pollution incidents’ throughout this article includes all incidents in categories 1-3. Where pollution incident severity is compared, all four pollution categories were used in the analysis. Notation is placed in the text when category 4 incidents are also included in analysis. Notation is also provided where sub-sets of categories are discussed.

EEA performance targets have evolved to reflect the ambition to continually reduce pollution incident numbers and increase self-reporting by the WASCs. This has led to periodic changes to EPA thresholds for the performance indicators in question. The EEA and Natural England (NE) outlined expectations for reduction of pollution incidents, in October 2017, within the ‘water industry strategic environmental requirements (WISER)’ report. The WISER document stated that “*serious pollution incidents (category 1 and 2) must continue to trend towards zero by 2020 with at least a 50% reduction compared to numbers of serious incidents recorded in 2012*” and that there should be a “*trend to minimise all pollution incidents (category 1-3) by 2025. There should be at least a 40% reduction compared to numbers of incidents recorded in 2016*”. Across WASCs there have been reductions in pollution incidents and increases in permit compliance and self-reporting since 2013 (EA, 2014; 2015; 2016b; 2017b; 2018; 2019; 2020). However, progress in pollution incident reduction and discharge permit compliance appears to have slowed, or declined for a number of WASCs, since 2016 (EA, 2017b; 2018; 2019; 2020). Due to continued pollution performance deterioration presented in EPA results over the last two years, the EEA have committed to a tougher regulatory approach (EA, 2019) with the introduction of Pollution Incident Reduction Plans required for all WASCs. Achieving a continued trend towards zero serious pollution incidents and 40% reduction in all pollution incidents by 2025, represents a significant challenge. In addition, WISER set out expectations for self-reporting; “*high levels of self-reporting of pollution incidents with at least 80% of incidents self-reported by 2025 and more than 90% of incidents self-reported for wastewater treatment works and pumping stations*”. Self-reporting has been assessed as above 80% for only four of the nine WASCs in the most recent EPA (EA, 2020b). In 2019, the chair of the EEA in England, Emma Howard Boyd, stated that “*rather than improving, the performance of most companies has deteriorated*” and in the 2020 EPA, stated that “*4 out of the 9 water companies are now rated as poor or requiring improvement, the worst result since 2011*” (EA, 2019; EA, 2020b).

Three out of the six performance indicators in the EPA are associated with pollution incidents (including pollution incident self-reporting by the WASCs). These performance indicators are calculated using data from the National Incident Reporting System (NIRS), which details environmental incidents within the remit of the EEA. Water and sewerage companies must record all potential and actual incidents on NIRS by their impact category (EA 2016b). The database holds a wealth of data on pollution incidents from WASCs, including the originating asset, whether it was self-reported and the impact to the water environment. The data are also used by the EEA in ongoing engagement with the WASCs. However, to the authors’ knowledge (and following a keyword search of academic journal outputs), there is no evidence to date of external peer-reviewed analysis and scrutiny of pollution incident performance data from the pollution incident databases across WASCs in England.

The EPA aims to provide a meaningful comparison of the variation in performance of the nine WASCs in England. Comparison of incident data from WASCs is made more difficult by the range of environmental and operational conditions the various WASCs function under. Previous authors in related disciplines have highlighted that operational variables (including company ownership, size, technology use, source of water, population density, energy consumption, construction year and the total area classed as rural) can affect efficiency levels in the water industry internationally (Conti, 2005; Abbott and Cohen, 2009; Munisamy, 2009; Renzetti and Dupont, 2009; Guerrini et al., 2011; Peda et al., 2011; Molinos-Senante et al., 2013; Walker et al., 2019). Thus, the operating environment of the WASCs could also impact pollution incident performance results calculated using the EPA performance indicator methodology. For example, pollution incident numbers are currently normalised per 10,000 km sewer length which is considered by the EEA the most user-friendly approach. This approach does not account for the number of total assets (including sewage treatment works, combined sewer overflows, and pumping stations) a company operates or the total population served. Deterioration in performance across the industry, makes it important to investigate common trends across all WASCs that could be addressed at a national and regional level. In addition, the findings of this research have implications for international water management, because trends in pollution performance could relate to declining water quality internationally (Chowdhury, 2018; Adams et al., 2019).

This research aims to analyse pollution incident data from WASCs in England, to assess and compare pollution incident performance in order to determine trends that could inform improvements to water management. The objectives of the study were; 1) to determine the differences in the pollution incident severity and sources across the WASCs; 2) to assess how the percentage of pollution incidents self-reported by the WASC affects the overall number of recorded pollution incidents; and 3) to determine if the number and length of asset types (operating property owned by the company, such as sewage treatment works and foul sewers) and population served influences pollution performance in each WASC.

## Methods

### Data Sources

The data analysed was obtained through Environmental Information Regulations 2004 requests to the EEA and includes;

- The National Incident Reporting System (NIRS) dataset 2010-2019 for the nine WASCs in England, including pollution incident data, location, offending asset type, pollution type, water impact and whether the incident was self-reported.
- WASC asset numbers and lengths (sewage treatment works, combined sewer overflows, pumping stations, foul sewers and rising mains) as used in the most recent EPA (EA, 2020b).
- WASC population served; as used in the most recent EPA (EA, 2020b).

### Performance indicator metric calculations

Within the EPA, current practice is to normalise pollution incidents by 10,000 km length of sewer. Sewer length, holds particular relevance to pollution incidents from the sewer network (primarily foul sewers and rising mains). Normalising pollution incident data by sewer length, may not take account of the variation evident in other sewage-related assets, seen in different WASCs operating areas (Table 1). For a more useful comparison of WASC pollution performance, the metric used to assess these companies, should account for variation in the WASC operating environment. Therefore, it was important to assess whether the method used to normalise data in current pollution incident EPA metrics impacted the performance assessment results. To assess the impact that the normalising variable has on the results of the EPA metrics, three alternative metrics normalised with different factors were investigated. It was hypothesised that variation in the results of current and alternative metric scores would be the consequence of variation in WASC operating environments (i.e. variation in the numbers of assets and the total sewer length).

**Table 1.**
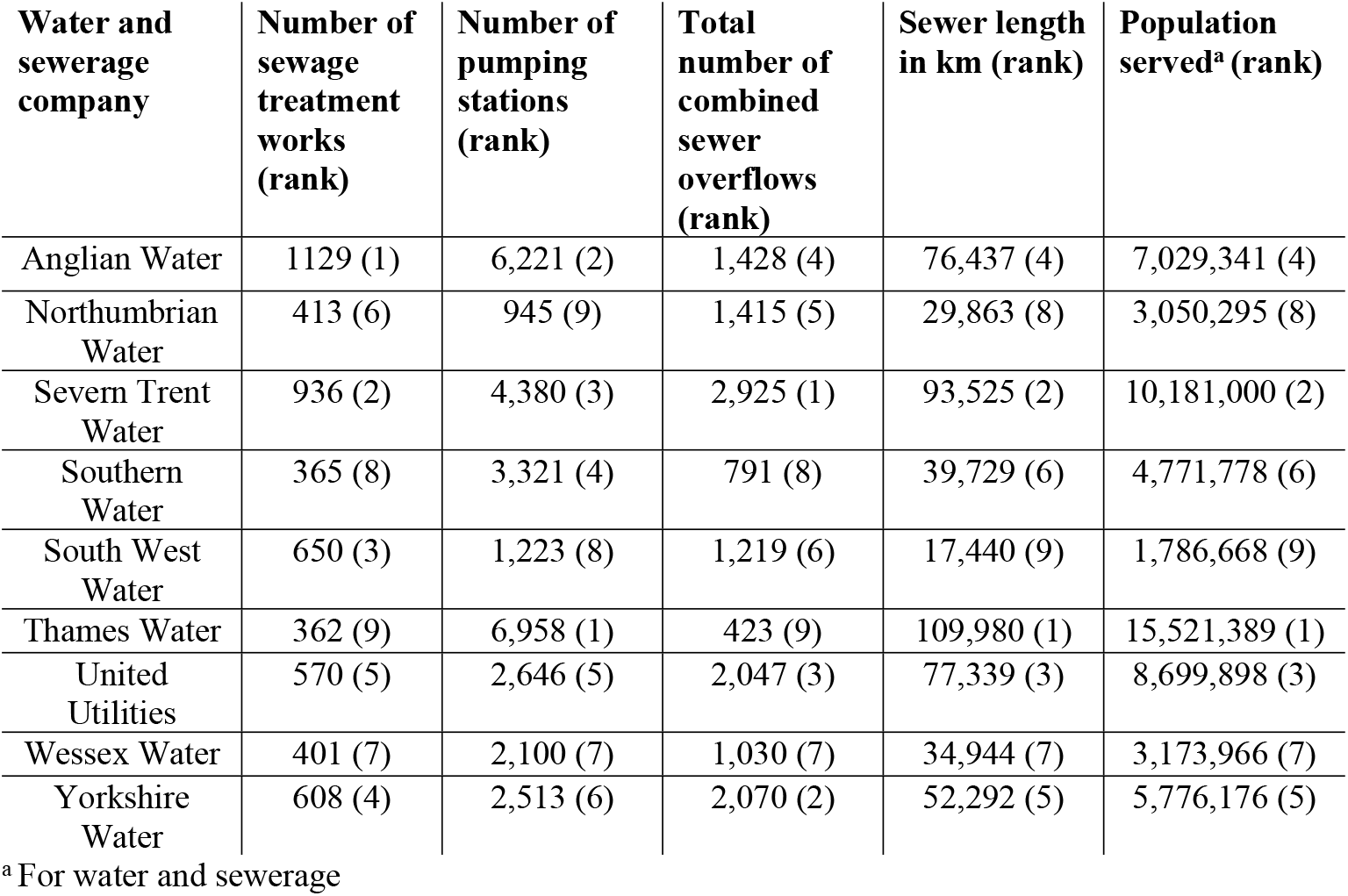
Asset numbers/lengths and population served for each water and sewerage company in 2020, with rank (1-9 in order of highest to lowest number/length)

| **Water and sewerage company** | **Number of sewage treatment works (rank)** | **Number of pumping stations (rank)** | **Total number of combined sewer overflows (rank)** | **Sewer length in km (rank)** | **Population served<sup>a</sup> (rank)** |
| --- | --- | --- | --- | --- | --- |
| Anglian Water | 1129 (1) | 6,221 (2) | 1,428 (4) | 76,437 (4) | 7,029,341 (4) |
| Northumbrian Water | 413 (6) | 945 (9) | 1,415 (5) | 29,863 (8) | 3,050,295 (8) |
| Severn Trent Water | 936 (2) | 4,380 (3) | 2,925 (1) | 93,525 (2) | 10,181,000 (2) |
| Southern Water | 365 (8) | 3,321 (4) | 791 (8) | 39,729 (6) | 4,771,778 (6) |
| South West Water | 650 (3) | 1,223 (8) | 1,219 (6) | 17,440 (9) | 1,786,668 (9) |
| Thames Water | 362 (9) | 6,958 (1) | 423 (9) | 109,980 (1) | 15,521,389 (1) |
| United Utilities | 570 (5) | 2,646 (5) | 2,047 (3) | 77,339 (3) | 8,699,898 (3) |
| Wessex Water | 401 (7) | 2,100 (7) | 1,030 (7) | 34,944 (7) | 3,173,966 (7) |
| Yorkshire Water | 608 (4) | 2,513 (6) | 2,070 (2) | 52,292 (5) | 5,776,176 (5) |

<sup>a</sup> For water and sewerage

The first alternative metric, normalised the total number of pollution incidents by total assets (per 100) for which a WASC is responsible for and included sewage treatment works, pumping stations and combined sewer overflow asset numbers. The second alternative metric, normalised total pollution incident numbers by each asset type and was calculated by dividing the number of pollution incidents emanating from each asset type by the number or length of that asset type. Pollution incidents from foul sewers and rising mains, were combined and normalised per 10,000km of total WASC sewer length. Pollution incidents from sewage treatment works, pumping stations and combined sewer overflows were normalised per 100 of these asset types operated by the WASC. Once calculated the normalised number for each asset type was combined to give an overall index score. Finally, the third alternative metric, normalised the total number of pollution incidents by the population served (per 100,000) in the WASC (water and sewerage) operating area.

### Statistical analysis

All data distributions were analysed for normality with the Anderson-Darling test, before further statistical analysis was conducted. Non-parametric tests were employed where data did not conform to the assumptions of parametric tests. Correlation analysis was performed using either the parametric Pearson’s or the non-parametric Spearman’s Rank correlation coefficients. Linear regression was used to determine the percentage of pollution incident variation that WASC self-reporting levels accounted for. The non-parametric Wilcoxon-Signed Rank test was used to determine if there were significant differences between paired data. All statistical tests were conducted using the software Minitab (Version 19) with a significance level set at p<0.05. The results of statistical tests are presented in parenthesis with the P value to support the interpretation within the text.

## Results and Discussion

### Comparison of water and sewerage company pollution incident severity and sources

Figure 1 displays pollution incident numbers by severity (category 1-4) from 2010-2019 for all nine WASCs in England. The large majority of pollution incidents reported were in the less serious categories 3 and 4, comprising between 97 and 99% of all pollution incidents reported in each year. An increase of category 3 and 4 pollution incident numbers from 2010 to a peak number in 2012 (2684 and 2265, in 2012, respectively) is evident. Numbers of category 3 and 4 pollution incidents decrease from 2012 to 2015. Personal communication with the EEA (2020), revealed that WASCs received pollution incident reporting training in 2014, because there was disparity in reporting practice, with some companies reporting more incidents than required. The incident report training appears to have reduced pollution incident reporting for category 3 and 4 pollution incidents in 2015. Subsequently, the number of category 3 and 4 pollution incidents have risen. There has been little variation in the more serious category 1 and 2 pollution incidents since 2014. Statistical analysis indicates that category 1-4 incidents were significantly greater in 2019 compared to 2018 (p =<0.001; Wilcoxon Signed-Rank Test).

**Fig 1:**
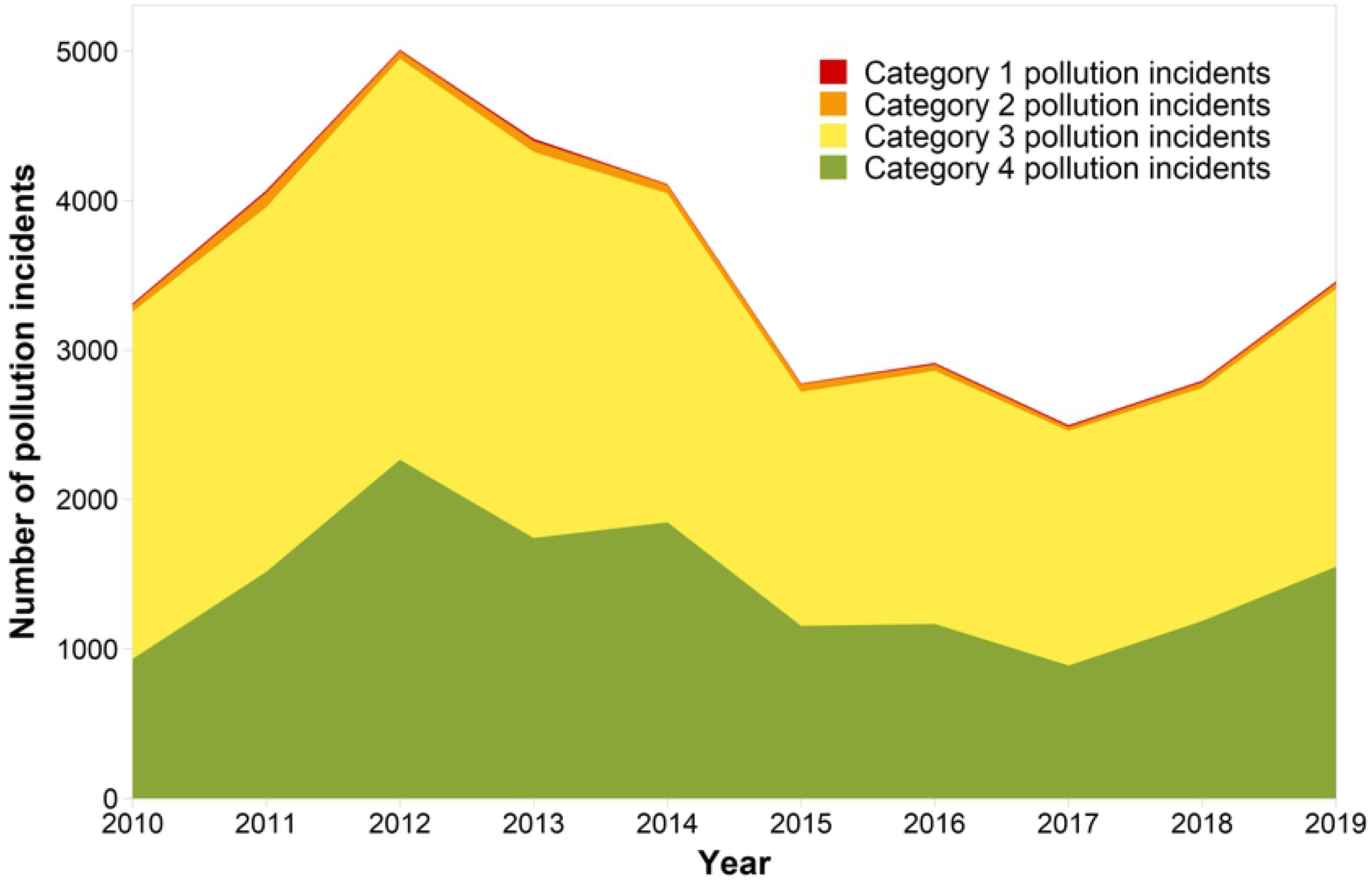
The total number of pollution incidents reported by all water and sewerage companies combined in England for pollution incident category 1, 2, 3 and 4 (2010-2019). Category 1 = major, serious, persistent and/or extensive impact or effect on the water environment. Category 2 = significant impact or effect on the water environment. Category 3 = minor or minimal impact or effect on the water environment. Category 4 = substantiated incident with no impact.

Figure 2 displays the number of pollution incidents impacting the water environment (categories 1-3) by asset type for all WASCs combined from 2010-2019. Key asset types where the majority of incidents took place are foul sewers, pumping stations, sewage treatment works, rising mains and combined sewer overflows. These asset types will be the focus of subsequent analysis. Foul sewers contributed the highest number of pollution incidents impacting the water environment from 2010-2019 (n=8515). Pollution incident numbers have remained relatively consistent from foul sewers annually since 2010 (mean = 857, standard deviation = 114). Annually, numbers of pollution incidents from rising mains have also remained relatively consistent and in relatively low numbers (mean = 121, standard deviation = 14). For numbers of pollution incidents from combined sewer overflows, there is a trend of reducing numbers from 2010 onwards (288 in 2010 to 78 in 2019). It is important to note that in some cases when other assets have faults, they may pollute through combined sewer overflows, but this would be recorded against the faulting asset. Numbers of pollution incidents have increased for sewage treatment works (from 321 in 2018 to 381 in 2019), but the highest increase in pollution incidents in 2019, was observed from pumping stations; an increase from 2018 levels of 170 (from 312 to 482). For numbers of all pollution incidents, pumping stations are the 2^nd^ most numerous source (n= 5022 from 2010-2019) and they represent the asset where there is greatest variation, followed by foul sewers and combined sewer overflows (Figure 2). For significant and major pollution incidents (category 1 and 2) declines are evident across all asset types in 2012 and 2017, but there have been no consistent reductions post 2014.

**Fig 2:**
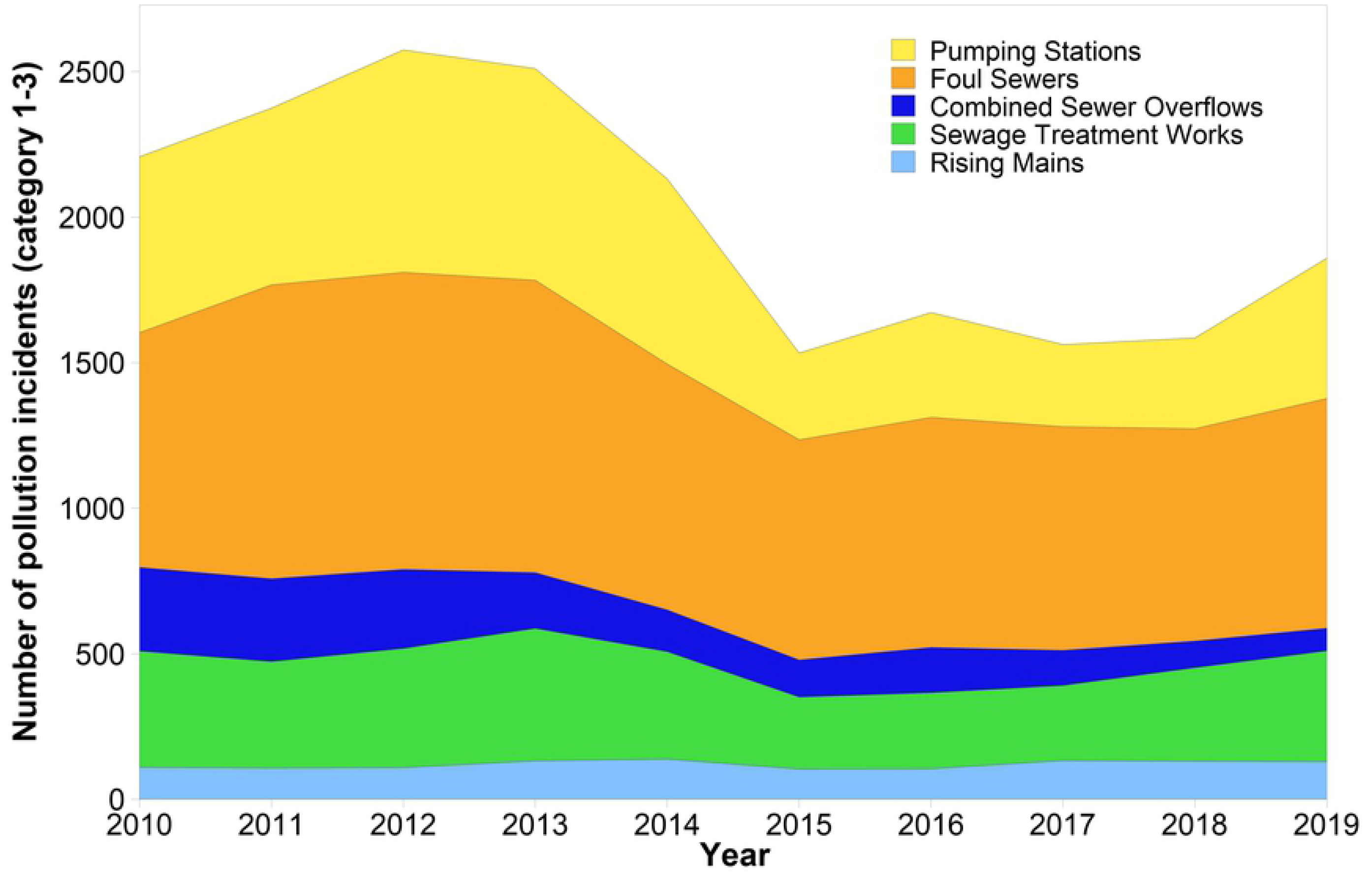
Number of pollution incidents for all water and sewerage companies combined in England for pollution category 1 to 3 by asset type (pumping stations, foul sewers, combined sewer overflows, sewage treatment works and rising mains) 2010-2019. Category 1-3 = pollution incidents with impact or effect on the water environment.

Figure 3 shows the number of pollution incidents by asset type and WASC. The companies are anonymised randomly and coded A-I. Figure 3, shows considerable variation in pollution incident numbers for different asset types across the individual WASCs. For most WASCs there is a decline in the number of pollution incidents recorded post 2014, when pollution incident training was implemented by the EEA, as described for the sector above. Companies B, D, E, G and H display increases in pollution incidents from 2015 onwards. It is notable that for company H, there is a substantial increase in numbers of pollution incidents in 2019 from pumping stations and sewage treatment works in comparison to the period from 2015-2018. This shift in the number of pollution incidents across these asset types indicates a potential change in reporting practise across the company or a significant decline in performance. From 2018 to 2019, WASC H, increased its self-reporting of pollution incidents from pumping stations from 88.1% to 97.5% and in sewage treatment works from 90.9% to 95.1%, respectively. WASCs D and E also show increases in the total numbers of pollution incidents originating from pumping stations and like company H, also increased the self-reporting of pollution incidents from this asset type to 90.0% and 92.4% from 72.7% and 81.1%, respectively. Conversely, companies C and I recorded fewer pollution incidents from pumping stations in 2019 and had decreases in self-reporting percentages of pollution incidents from this asset type (6.5% and 6.6%, respectively). These trends were not evident for all companies, with company F increasing the self-reporting percentages for pollution incidents originating from pumping stations from 42.9% to 80.0%, whilst also recording fewer total pollution incident numbers from this asset type. However, it is important to highlight that the percentages calculated for company F are based on a low number of pollution incidents from this asset type (7 in 2018 and 5 in 2019). Similar trends can be observed for sewage treatment works. Companies F, G, and H all had increased pollution incident self-reporting percentages for sewage treatment works (between 2 and 9%) and increases in pollution incident numbers (between 7 and 69). Companies A, C and I, recorded decreases in pollution incident self-reporting percentage (between 7 and 67%) and pollution incident number reductions of between 8 and 26. Only company D managed to simultaneously increase self-reporting (up by 7%) and decrease pollution incident numbers from sewage treatment works (down by 1). These results suggest that increases in pollution incidents from pumping stations and also sewage treatment works are largely responsible for declines in WASC performance in EPA pollution incident metrics. Observation of these trends make it important to determine the influence self-reporting has on pollution incident numbers from individual asset types. The authors hypothesised that increases in self-reporting performance could be leading to the recording of previously uncaptured pollution incidents from some asset types.

**Fig 3:**
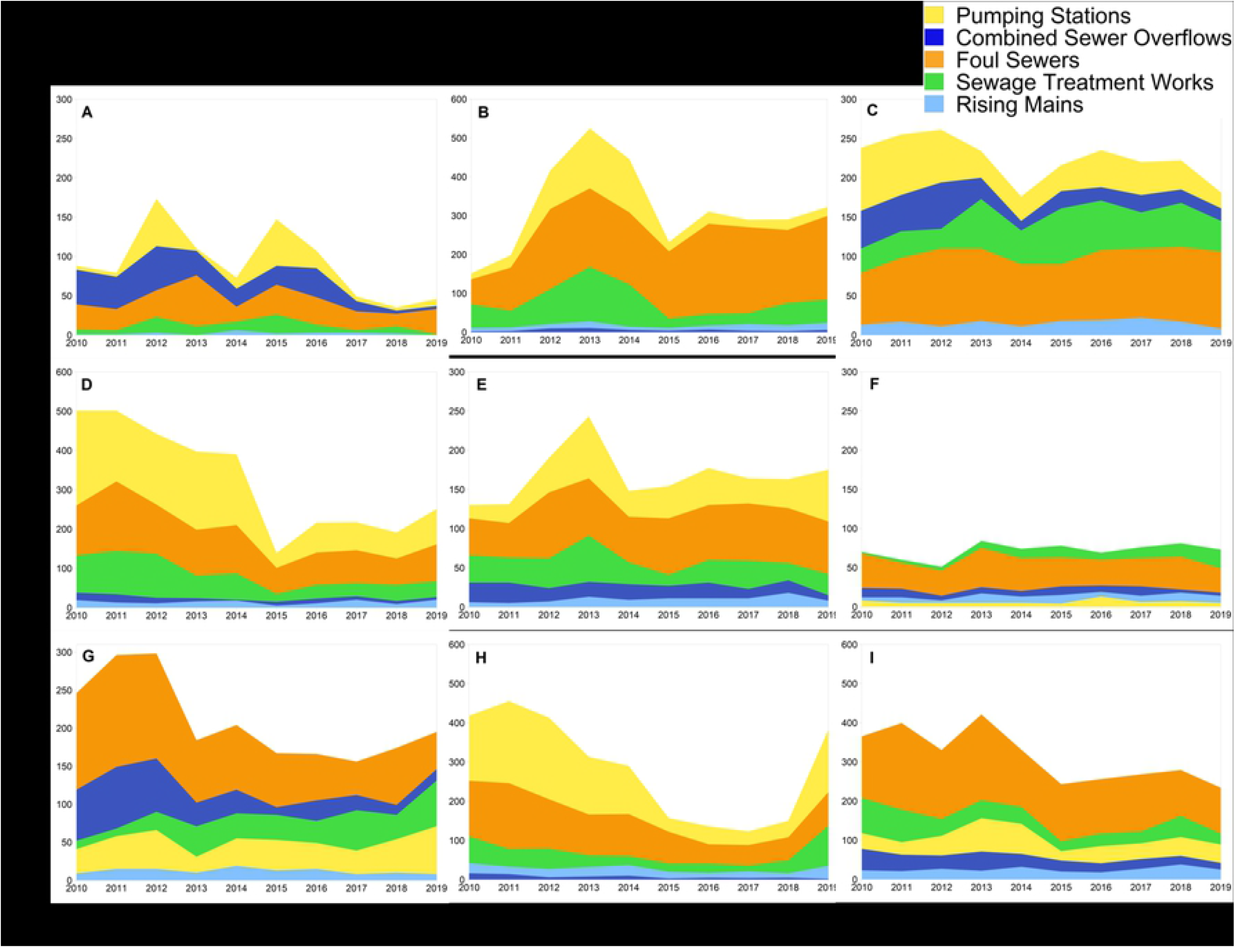
Number of pollution incidents by water and sewerage company (anonymised and coded A-I) in England for pollution categories 1 to 3 by asset type (pumping stations, foul sewers, combined sewer overflows, sewage treatment works and rising mains (2010-2019). Category 1-3 = pollution incidents with impact or effect on the water environment.

### Water and sewerage company self-reporting

Figure 4 displays the percentage of pollution incidents self-reported by WASCs in England by the asset type for 2019. Whilst self-reporting across all assets has improved from 2018 (EA, 2019), six companies self-reported less than 80% of pollution incidents. Self-reporting percentages are not consistent across asset types, even for companies performing well in this EPA indicator. The largest range in self-reporting percentages across the WASCs is shown for combined sewer overflows (between 16% and 100%). Lower self-reported pollution incident percentages are also evident for foul sewers (between 50% and 82%). This might be because pollution incidents from foul sewers tend to be quickly seen by members of the public and it is more obvious when an incident is occurring in these settings. However, for asset types more commonly associated with remote or less populated areas (such as sewage treatment works, combined sewer overflows and pumping stations), it may be less likely that a member of the public would report a pollution incident and/or consider a discharge from these settings to be unusual, unless the incident is of a more significant nature. Figure 5 displays the number of pollution incidents (category 1-4) that were self-reported and not self-reported by the WASCs combined (in 2019) by all five asset types. A greater proportion of the pollution incidents not self-reported by the WASCs have a more serious impact (category 1-2). This suggests that slower reaction times to incidents that are not self-reported could increase the likely impact to the water environment and/or that pollution incidents with larger impact are more likely to be spotted and reported by non-WASC sources. The results indicate that lower impact pollution incidents (category 3 and 4), that are not self-reported, may go unreported or progress into more serious incidents (category 1 and 2) based on higher proportions of category 1 and 2, and lower proportions of category 3 and 4 pollution incidents that were reported by non-WASC sources.

**Fig 4:**
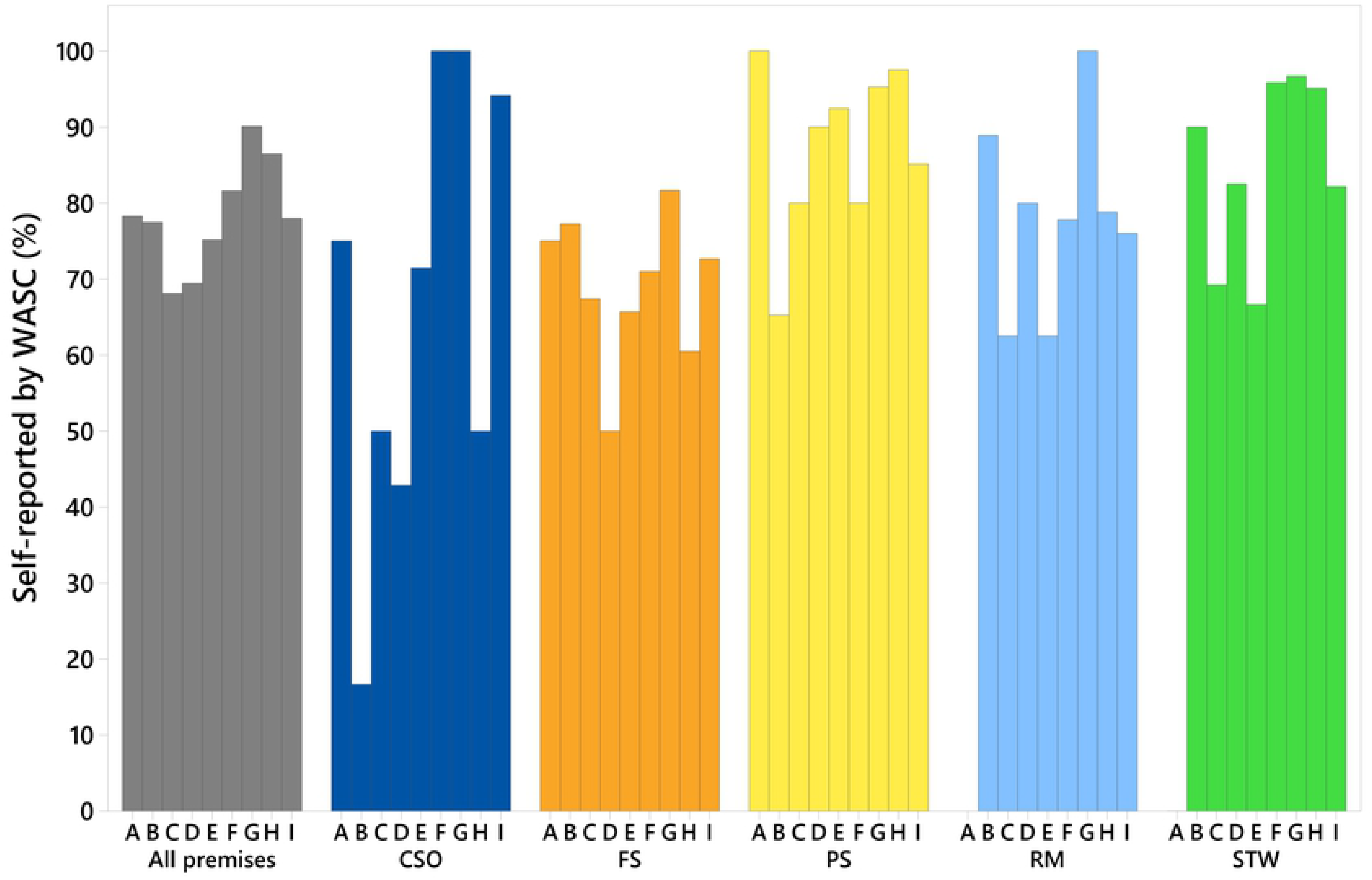
Percentage of pollution incidents (category 1-3) self-reported by water and sewerage companies (anonymised and coded A-I) in England by the asset types; all assets (premises), combined sewer overflows (CSO), foul sewers (FS), pumping stations (PS), rising mains (RM) and sewage treatment works (STW) (2019). Category 1-3 = pollution incidents with impact or effect on the water environment.

**Fig 5:**
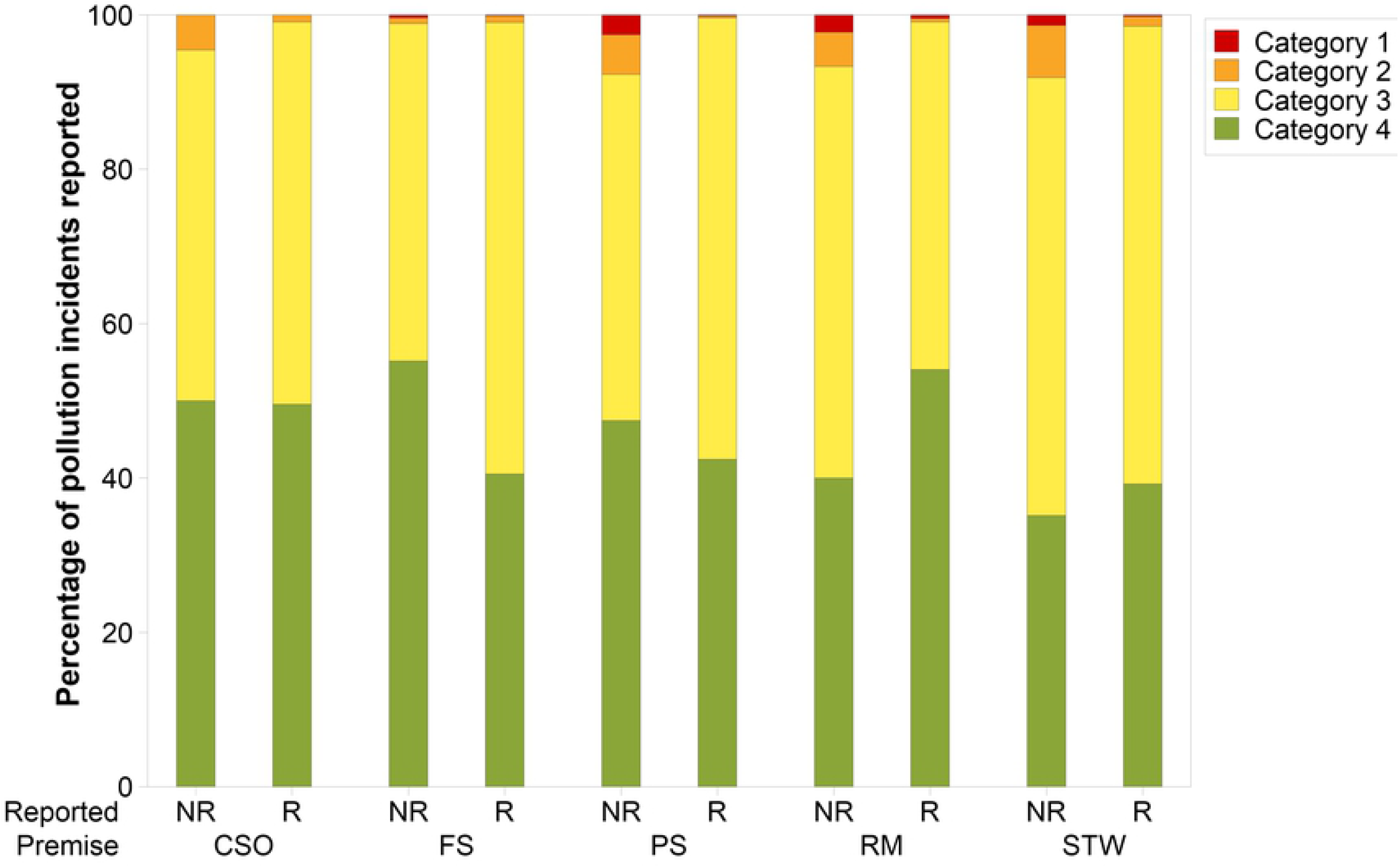
Percentage of pollution incidents (category 1-4) that were self-reported (R) and not self-reported (NR) by the water and sewerage companies combined in 2019 by the asset types; combined sewer overflows (CSO), foul sewers (FS), pumping stations (PS), rising mains (RM) and sewage treatment works (STW). Category 1 = major, serious, persistent and/or extensive impact or effect on the water environment. Category 2 = significant impact or effect on the water environment. Category 3 = minor or minimal impact or effect on the water environment. Category 4 = substantiated incident with no impact.

Variation in self-reporting across the asset types for different WASCs, observed in Figure 4, warrants further investigation. Variation could impact the pollution incident numbers reported, if low self-reporting percentages result in incidents not being reported. To determine if self-reporting percentages for asset types have an influence on the pollution incident performance of the WASCs, correlation and regression analysis was performed. Relationships between the number of pollution incidents normalised by 10,000 km of sewer length and the percentage of incidents that were self-reported by the WASCs in England, were investigated using data from 2010-2019 (n=90 for each test) (Figure 6). Spearman’s Rank correlation analysis, revealed statistically significant positive relationships between pollution incident self-reporting percentages and numbers of pollution incidents per 10,000 km sewer length for pumping stations and sewage treatment works (r=0.215, P-value= 0.042 and r= 0.245 and P-value= 0.020, respectively). Statistically significant relationships were not evident between the pollution self-reporting percentage and numbers of pollution incidents for other asset types investigated. Regression analysis indicated that self-reporting percentages explained a low percentage of the variation in pollution incidents per 10,000 km from 2010 to 2019 for pumping stations (R-sq = 4.9 %)) and sewage treatment works (R-sq = 5.2 %). The results do indicate that for pumping stations and sewage treatment works, a number of pollution incidents may be going unreported when self-reporting percentages are lower than 100 %.

**Fig 6:**
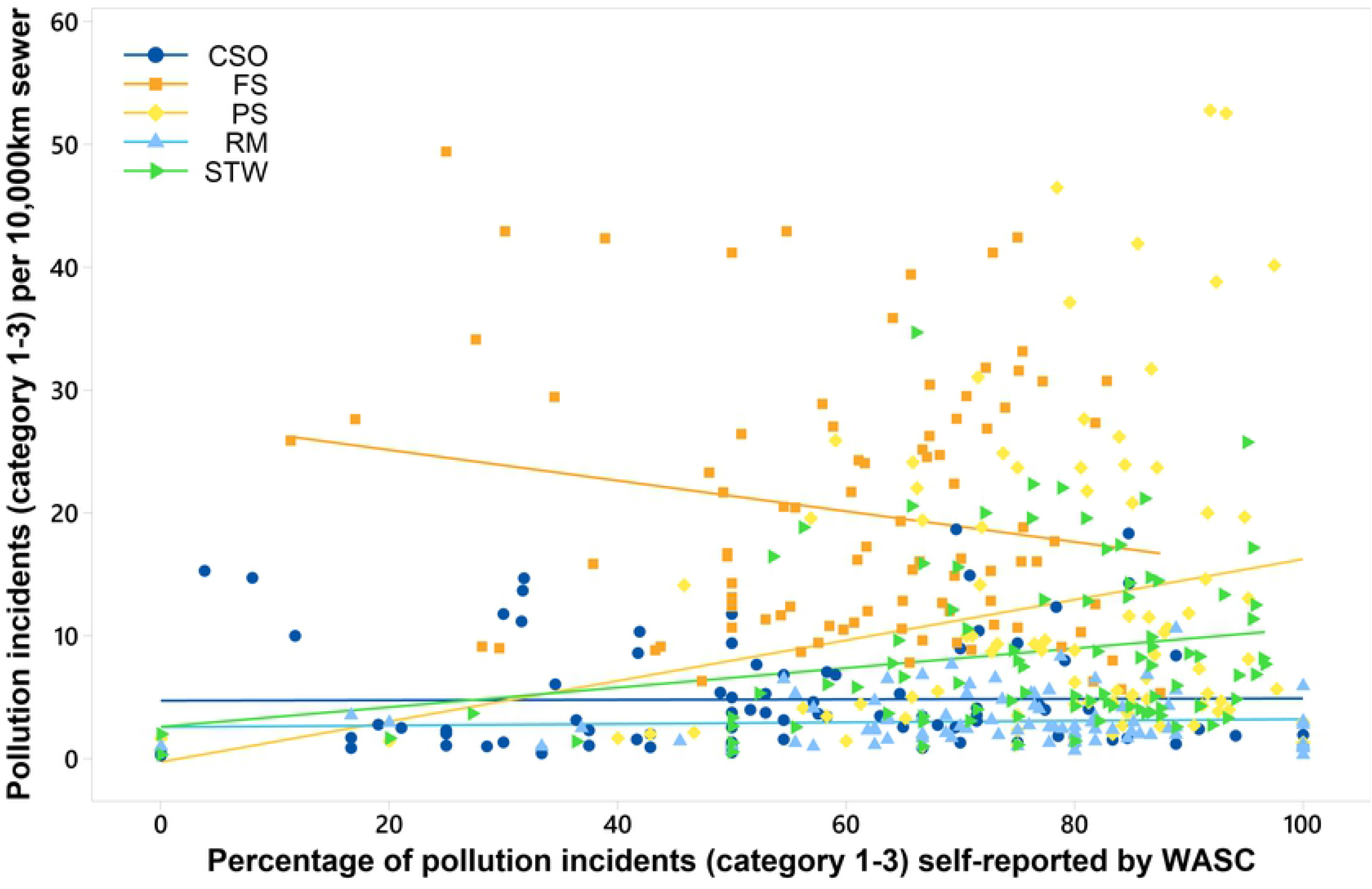
Relationships between pollution incidents (categories 1-3) per 10,000 km sewer length and the percentage of pollution incidents (categories 1-3) self-reported by all water and sewerage companies in England (2010-2019) according to asset type: combined sewer overflows (CSO), foul sewers (FS), pumping stations (PS), rising mains (RM), and sewage treatment works (STW). Lines represent the best regression fit. n=90 for each asset type. Category 1-3 = pollution incidents with impact or effect on the environment, people and/or property.

Correlation analysis was used to assess independent relationships between the numbers of pollution incidents per 10,000km sewer length from individual asset types and the self-reporting percentage for that asset for each WASC. Note that these results should be interpreted with caution as there are relatively few data (n=10 for each WASC). Correlation analysis between numbers of pollution incidents per 10,000km and self-reporting percentages of particular asset types revealed that for some WASCs, correlation was significant and much stronger for individual companies, then was observed for WASC data combined (Table 2).

**Table 2:** Statistically significant correlations between numbers of pollution incidents (category 1-3; pollution incidents with impact or effect on the water environment) per 10,000km sewer length and self-reporting percentages for asset types by water and sewerage company (n=10 for each water and sewerage company).

| Water and sewerage company code | Asset type | Correlation coefficient | P-value | r |
| --- | --- | --- | --- | --- |
| B | Foul sewers | Spearman's Rank | 0.019 | 0.721 |
| C | Combined sewer overflows | Spearman's Rank | 0.003 | 0.832 |
|  | Foul sewers | Pearson's | 0.038 | 0.661 |
|  | Rising mains | Pearson's | 0.011 | 0.758 |
| D | Pumping stations | Pearson's | 0.037 | 0.661 |
| E | Rising mains | Pearson's | 0.003 | 0.824 |
| F | Sewage treatment works | Pearson's | 0.047 | 0.638 |
| G | Sewage treatment works | Spearman's Rank | 0.004 | 0.815 |
| H | Foul sewers | Pearson's | 0.027 | 0.692 |
|  | Sewage treatment works | Spearman's Rank | 0.028 | 0.687 |
|  | Rising mains | Pearson's | 0.014 | -0.740 |
| I | Combined sewer overflows | Pearson's | 0.000 | -0.908 |

While smaller data volumes could inflate correlation strengths, these results indicate that for some WASCs, increases in self-reporting could lead to a greater number of total pollution incidents recorded. The strongest significant positive relationship was observed for WASC C and combined sewer overflows (P-value= 0.003 and r= 0.832). Company C, has improved self-reporting of pollution incidents from combined sewer overflows, but self-reporting percentages remain low at 50%. Therefore, correlation could suggest that increased self-reporting in this asset type for company C is leading to increased pollution incident numbers, because of the capture of incidents that would have previously gone unreported. In support of this theory, recent research conducted by Hammond et al. (2021) reported 926 unreported putative ‘spills’ from combined sewer overflows as determined by machine learning techniques, in only two wastewater treatment works in England. Similarly, strong significant positive relationships were evident for sewage treatment works (r = 0.815) and rising mains (r = 0.824) for company G and E, respectively. Negative correlations were also observed, whereby self-reporting has increased in an asset type, along with decreasing pollution incidents. For company H and I, significant strong negative relationships were apparent for the number of pollution incidents per 10,000 km sewer length and self-reporting percentages from rising mains (company H, r = -0.740) and combined sewer overflows (company I, r = - 0.908). In these instances, the authors suggest that this is reflective of increased performance with decreased numbers of pollution incidents in these asset types as a result. Whilst a number of other variables could have also led to an increase in pollution incidents for these WASCs, including extreme meteorological conditions, aging assets and increases in populations served; these results do indicate that self-reporting percentage could be an important contributor to variation.

Table 3 displays the number of water company reported pollution incidents between 2010 and 2019 for all asset types investigated. The number and percentage of the water company pollution incidents that were also reported by a non-water company source (duplicate report) are also given. For all asset groups investigated 7% or less were also reported by a non-water company entity. Duplicate reports from non-water company entities may indicate that the incident would have been reported even if the water company did not self-report the incident. Some incidents may not receive duplicate reports, because action on site to resolve the incident may make it obvious a report has been received already. However, a lack of duplicate report, may also indicate that incidents not self-reported are unlikely to be reported by non-water company entities.

**Table 3:** Water company reported pollution incidents (category 1-3; pollution incidents with impact or effect on the water environment) between 2010 and 2019 for combined sewer overflows, foul sewers, pumping stations, rising mains and sewage treatment works and the number and percentage of these incidents where a non-water company source duplicated the report

| Asset type | Pollution incidents reported by water companies 2010-2019 | Number of pollution incidents also reported by a non-water company source 2010- 2019 (percentage is shown in brackets) |
| --- | --- | --- |
| Combined Sewer Overflow | 1018 | 35 (3%) |
| Foul Sewer | 5334 | 288 (5%) |
| Pumping Station | 4134 | 115 (3%) |
| Rising Main | 921 | 65 (7%) |
| Sewage Treatment Works | 2801 | 49 (2%) |

These results highlight the importance of robust reporting standards and positive reporting culture, which could include a shift towards higher self-reporting targets. If pollution incidents are being missed, as this research indicates, WASCs should be aiming to self-report all pollution incidents for all asset types, especially pumping stations and sewage treatment works. These results suggest that current self-reporting performance metrics need to include a breakdown of self-reporting percentages across all asset types.

### Water and sewerage company pollution performance

To determine if the number and/or length of asset types, and population served, influences pollution performance in each WASC, alternative pollution performance metrics from those currently employed in the EPA were assessed. Table 1 displays the variation between WASCs in asset numbers and lengths, as well as the populations served in relation to sewer length (km). This variation prompts investigation into the most suitable metric for standardised comparison across all WASCs. Table 4 presents the results of three alternative pollution performance metrics, in comparison to the current EPA metric for total pollution incidents. The rank of the WASC according to the results of each alternative metric are shown, along with the difference to the initial rank, which was calculated using EPA metric methodology. The results show that for the highest performing company (A), changes in the pollution performance metric make no difference to the overall ranking. This would suggest that the performance would be considered industry leading, regardless of the company operating conditions. Similarly, for company F, alternative metric 1 and 2, have no effect on the 2^nd^ place ranking. Of the metric results, variation from the original ranking is greatest when normalising total pollution incidents per 100,000 population served (alternative metric 3). For a standardised and consistent comparison of WASC pollution performance, a metric based on population served may be difficult to implement, because some WASCs may receive larger volumes of visitors throughout the year with peaks in summer months more likely. Whilst there is variation between the companies initially ranked 8 and 9, they remain in the bottom three for all metrics investigated. In general, more variation is evident for companies with an initial rank of between 3 and 7. This suggests that variation in the WASC operating environment could lead to differences in assessed pollution performance, for different pollution performance metrics for some WASCs. Whilst differences in rank are limited to ±2 for alternative metrics 1 and 2, the choice of metric for standardised comparison is important and should be considered carefully. Alternative metric 2 considers the largest proportion of asset variation by normalising pollution incidents by each asset type and then combining these scores. This metric or variations of it may provide the most reliable comparison of WASCs numerically. Another consideration is the ease of interpretation and how meaningful results are to the public. In this respect, the current metric using sewer length is still clearer for non-specialist audiences, and then alternative metric 2. Currently, these scores are not assessed with thresholds, such as those defined in the current EPA (EA, 2017a). Rankings are not employed in the EPA, and whilst useful for assessing differences for the purpose of this study, it will be important to also assess differences in threshold assignments in consultation with the EEA.

**Table 4:**
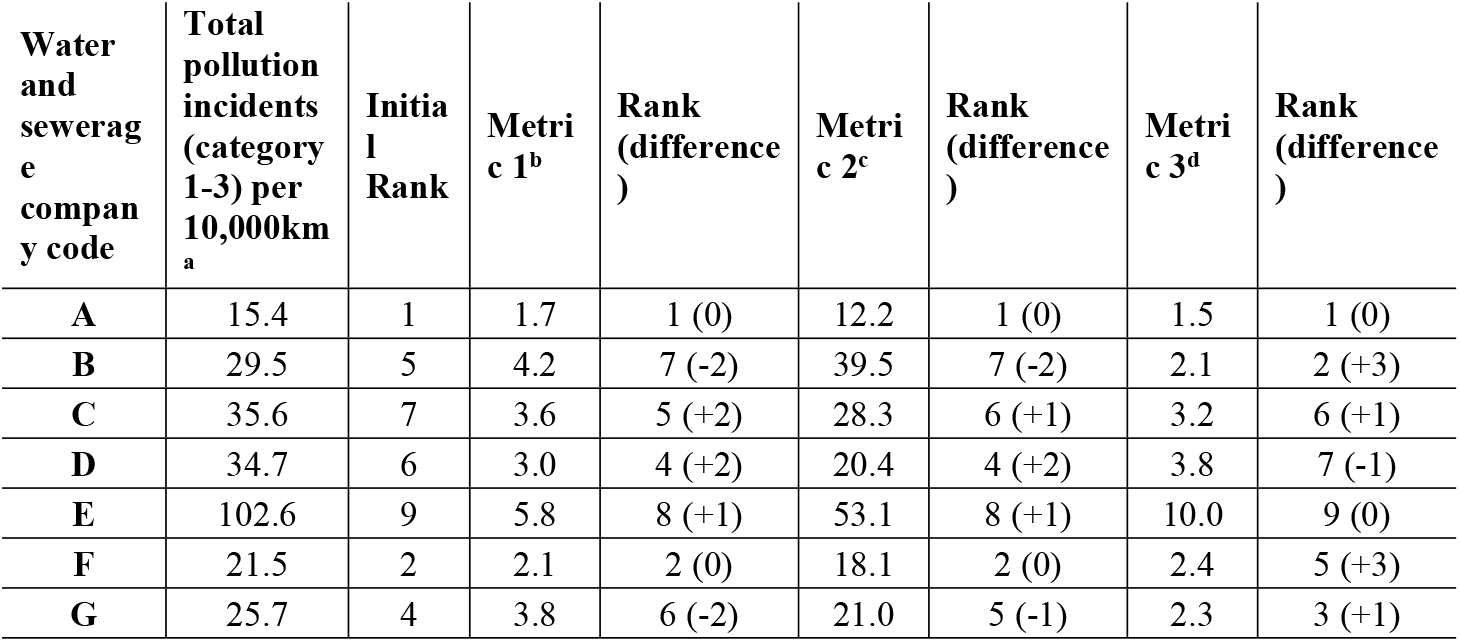
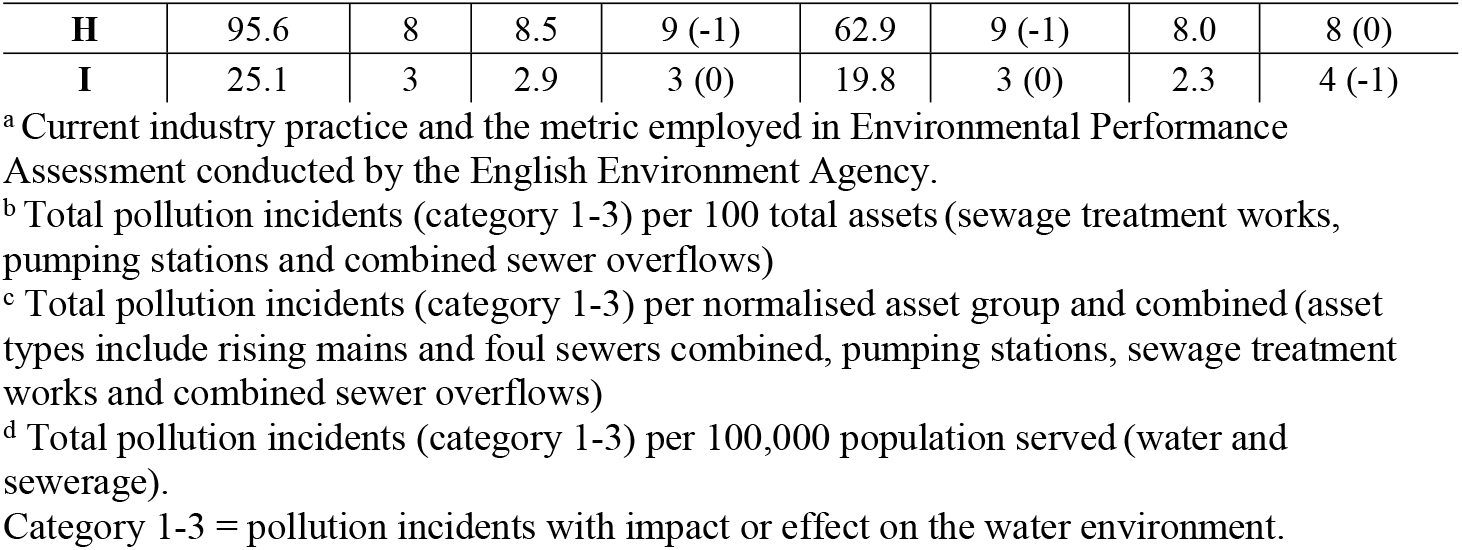
Water and sewerage company pollution performance in 2019 according to pollution incidents causing a negative impact to the water environment, normalised by 1) sewer length, total assets, 3) each asset type, and 4) the population served

| Water and sewerage company code | Total pollution incidents (category 1-3) per 10,000km<sup>a</sup> | Initial Rank | Metric 1<sup>b</sup> | Rank (difference) | Metric 2<sup>c</sup> | Rank (difference) | Metric 3<sup>d</sup> | Rank (difference) |
| --- | --- | --- | --- | --- | --- | --- | --- | --- |
| A | 15.4 | 1 | 1.7 | 1 (0) | 12.2 | 1 (0) | 1.5 | 1 (0) |
| B | 29.5 | 5 | 4.2 | 7 (-2) | 39.5 | 7 (-2) | 2.1 | 2 (+3) |
| C | 35.6 | 7 | 3.6 | 5 (+2) | 28.3 | 6 (+1) | 3.2 | 6 (+1) |
| D | 34.7 | 6 | 3.0 | 4 (+2) | 20.4 | 4 (+2) | 3.8 | 7 (-1) |
| E | 102.6 | 9 | 5.8 | 8 (+1) | 53.1 | 8 (+1) | 10.0 | 9 (0) |
| F | 21.5 | 2 | 2.1 | 2 (0) | 18.1 | 2 (0) | 2.4 | 5 (+3) |
| G | 25.7 | 4 | 3.8 | 6 (-2) | 21.0 | 5 (-1) | 2.3 | 3 (+1) |
| **H** | 95.6 | 8 | 8.5 | 9 (-1) | 62.9 | 9 (-1) | 8.0 | 8 (0) |
| **I** | 25.1 | 3 | 2.9 | 3 (0) | 19.8 | 3 (0) | 2.3 | 4 (-1) |

<sup>a</sup> Current industry practice and the metric employed in Environmental Performance Assessment conducted by the English Environment Agency.
<sup>b</sup> Total pollution incidents (category 1-3) per 100 total assets (sewage treatment works, pumping stations and combined sewer overflows)
<sup>c</sup> Total pollution incidents (category 1-3) per normalised asset group and combined (asset types include rising mains and foul sewers combined, pumping stations, sewage treatment works and combined sewer overflows)
<sup>d</sup> Total pollution incidents (category 1-3) per 100,000 population served (water and sewerage).
Category 1-3 = pollution incidents with impact or effect on the water environment.

Companies are not required to report category 4 pollution incidents (substantiated incident with no impact). Therefore, category 4 incidents recorded in the NIRS database, are likely to be the result of WASCs challenging a higher impact category and successfully downgrading the impact level. Thus, differences noted in calculated ratios of category 1-3 to category 4 pollution incidents for different WASCs (0.86-1.75) may reflect more frequent or more successful challenges of impact classification or vice versa. Category 4 data are not currently checked by the EEA and as a result the data may not have the same reliability as recorded category 1-3 pollution incidents. Checks and assurance on category 4 data could enable further assessment of pollution incident challenge influence on performance in the future.

### Pollution incident impacts

Pollution incidents have the potential to be detrimental to ecosystems and human health. In worst case scenarios pollution incidents can result in large scale aquatic organism mortality events. Research has also suggested that frequent low impact events, that lead to long-term exposure to contaminants can be just as damaging to aquatic organisms (Saaristo et al., 2018). In one example, Mallin et al. (2007) investigated the impact of a raw sewage spill on water and sediment quality in an urbanised estuary in North Carolina, USA. Increased biochemical oxygen demand led to a large fish kill and high concentrations of nutrients also led to several algal blooms. Faecal bacteria levels were elevated in the water column and in the sediment. Faecal bacteria in the sediment were shown to persist, much longer than in the water column, indicating the long-term potential for contaminant storage in environmental reservoirs and resuspension in sediments downstream from sewage spills. Of particular concern is the potential for antibiotic resistance to spread in the environment, where antibiotics, antibiotic resistant bacteria, and antibiotic resistant genes enter water and sediments (Baquero et al., 2008). Another concern, is the introduction of human pathogens into surface waters used for recreation (Purnell et al., 2020). In addition, incidents can be a source of toxic metals and engineered particles (Emmons et al., 2018; Loosli et al., 2019). These are relatively few published examples of the serious implications of sewage spills to the environment but it is clear from the literature that this is a global issue. In light of the serious implications of pollution incidents as described above, the findings of this study are concerning. In England, it is clear there is an urgent need to reverse a trend of declining pollution performance across the water and sewerage sector. Immediate in-depth and company specific assessments of the poorly performing assets, primarily pumping stations and sewage treatment works, is recommended. The new requirement for WASCs to produce Pollution Incident Reduction Plans, including detailed analysis of pollution incident causes and strategies for improvements, is an important first step in improving pollution incident performance nationally. Increased self-reporting of pollution incidents may be inflating numbers, but it is clear that other factors are contributing to the decline in performance. These might include but are not limited to aging assets and infrastructure, changing climate and increasing populations. It was beyond the scope of this study and the data collected to investigate these factors, but future research is required to determine their influence on pollution incident frequency and severity.

## Conclusions

Whilst this research highlighted a number of key areas for more detailed analysis, in the short-term, research should focus on investigating best practice for reporting pollution incidents. It is important to get an accurate baseline of the number of pollution incidents and whether a proportion are currently going unreported. This is also vital for a fair comparison of WASC performance across England. This research suggests that events that are not reported quickly by the WASC, can lead to more severe impacts to the water environment. So rapid and consistent reporting of incidents is crucial for limiting damage. The results indicate that current self-reporting performance metrics need to include a breakdown of self-reporting percentages across all asset types. In addition, this research suggests that a focus on improvements to pumping stations and sewage treatment works in England in the first instance, would be prudent to reverse the declining pollution performance across the water and sewerage sector.

## Acknowledgements

The authors thank Southern Water Services (UK) for funding this research. The authors would also like to thank the English Environment Agency for providing data and personal communication throughout the length of this project.

